# Characterization of antibiotic resistance in clinical isolates of *Klebsiella pneumoniae* in Denmark

**DOI:** 10.1101/2021.12.20.473592

**Authors:** Xin Fang, Henrik Westh, Michael Kemp, Svend Ellermann-Eriksen, Bernhard O. Palsson, Helle Krogh Johansen, Jonathan M. Monk

**Author notes:** **Correspondence** To whom correspondence should be addressed: Jonathan M. Monk, University of California, San Diego, 9500 Gilman Drive, La Jolla, CA 92093, Helle Krogh Johansen, Department of Clinical Microbiology, Rigshospitalet, Section 9301, Henrik Harpestrengs Vej 4A, 2100-Copenhagen Ø, Denmark.

## Abstract

*Klebsiella pneumoniae* (KP) is a major global health problem as it leads to hospital outbreaks all over the world and is becoming more difficult to treat due to its increasing antimicrobial resistance (AMR). Optimization and development of new treatments of KP requires understanding of its population structure and AMR properties. Therefore, in this study, we collected and sequenced 491 KP strains from four major Danish microbiology departments covering 51% of the Danish population. The isolates were whole genome sequenced (WGS), phenotypically characterized and compared with 2,124 KP strains from 13 different countries (PATRIC strains). We found that while genomic content varies significantly across the Danish strains, they also differ significantly from strains from other countries, due to the lack of certain AMR sequence types (e.g. ST258 and ST307) in Denmark. Genomic and experimental analysis suggest that Danish strains contain fewer virulence mechanisms and are more susceptible to antimicrobials compared to strains from other countries, likely due to the relatively low antibiotic usage in Denmark where 70% of hospital antibiotic usage is penicillins. We also identified potential novel AMR determinants to tigecycline through statistical analysis of genomic and phenotypic data. To conclude, we obtained a more comprehensive understanding of the KP strains in Denmark and provided valuable insights for future experiments and strategies to combat AMR in KP.

## INTRODUCTION

The natural reservoir for *Klebsiella pneumoniae* (KP) is the environment and KP is frequently found in water, sewage, soil and plant surfaces. KP has also developed into an opportunistic pathogen that causes serious hospital-acquired infections in humans, especially in neonates, the elderly and immunocompromised hosts (Podschun and Ullmann 1998). It is a significant threat to global public health and recognized as an ESKAPE pathogen (*Enterococcus faecium, Staphylococcus aureus, Klebsiella pneumoniae, Acinetobacter baumannii, Pseudomonas aeruginosa*, and Enterobacter spp.)(De Oliveira et al. 2020) due its capability to carry multiple antibiotic resistance genes (Wyres and Holt 2016).

Genomic content is highly variable across different KP strains. Three distinct species exist within KP: *K. pneumoniae* (KpI), *K*.*quasipneumoniae* (KpII) and *K. variicola* (Kp III) and KpII and Kp III are known to be less virulent (Long, Linson, et al. 2017). These species can be distinguished by 3-4% nucleotide divergence from the core genome (Wyres and Holt 2016). MALDI-TOF mass spectrometry was also shown to be able to precisely differentiate the KP complex members (Rodrigues et al. 2018). Previous studies have shown that KP has a large accessory genome, suggesting each strain could have distinct metabolic, virulent and antimicrobial resistance (AMR) capabilities (Holt et al. 2015). Various virulence factors contribute to their pathogenicity in humans, including their capsular polysaccharide, irons acquisition system and adherence factors (Clegg and Murphy 2016; Martin et al. 2018). Specifically, strains belonging to certain clonal groups have been widely associated with hospital outbreaks, including strains from ST258 and ST307 (Long, Wesley Long, et al. 2017).

KP is intrinsically resistant to ampicillin since the SHV beta-lactamase is a part of its core genome. In addition, KP strains often acquire AMR gene through horizontal gene transfer via plasmids and more than 100 different AMR determinants have been identified in KP (Holt et al. 2015) that may confer resistance to almost all available antibiotics. Specifically, genes encoding a carbapenemase that can hydrolyze carbapenems (David et al. 2019; Munoz-Price et al. 2013) and genes encoding extended spectrum beta-lactamase (ESBL) are of particular concern in KP.

The treatment of KP has become increasingly difficult due to the aquisition of AMR determinants. To develop strategies to combat AMR, it is important to understand their population structure and AMR properties. Therefore, efforts in collecting, sequencing and characterizing KP strains have been made across the world (Moradigaravand et al. 2017; Henson et al. 2017; Loraine et al. 2018). In this study, we collected a set of 491 strains from four major Danish hospitals and characterized their genome content and antibiotics resistance capabilities. We have obtained a better understanding of KP strains in Denmark and predicted potential new AMR determinants to be validated by experiments.

## RESULTS

### Collection of Danish Strains

We have collected 491 *Klebsiella Pneumoniae* (KP) blood culture isolates from patients from four Danish clinical microbiology departments with an uptake area covering 51% of the Danish population between January 2018 to December 2018. Of all 491 strains, 38.1% (187) were from Aarhus University Hospital (AUH), 30.3% (149) from Hvidovre Hospital (HVH), 19.6% (96) from Odense University Hospital (OUH), and 12.0% (59) from Rigshospitalet (RH), which is a tertiary referral hospital and the most specialised hospital in Denmark(Fig. 1A). These strains were isolated from 165 female patients and 326 male patients between 4 to 104 years of age, and the majority of the strains were isolated from patients between 64-84 years of age (Fig. 1B). Of all strains collected, 27.4% came from the Emergency department, and the rest of the strains were collected in Urology, Intensive Care Units, Nephrology and various other clinical departments (Fig. 1C). Two microbiology departments did not submit data on clinical departmental information for 52.1% of isolates. Both genome sequences and antimicrobial resistant profiles were generated for these strains and will be discussed in detail in the following sections.

**Figure 1:**
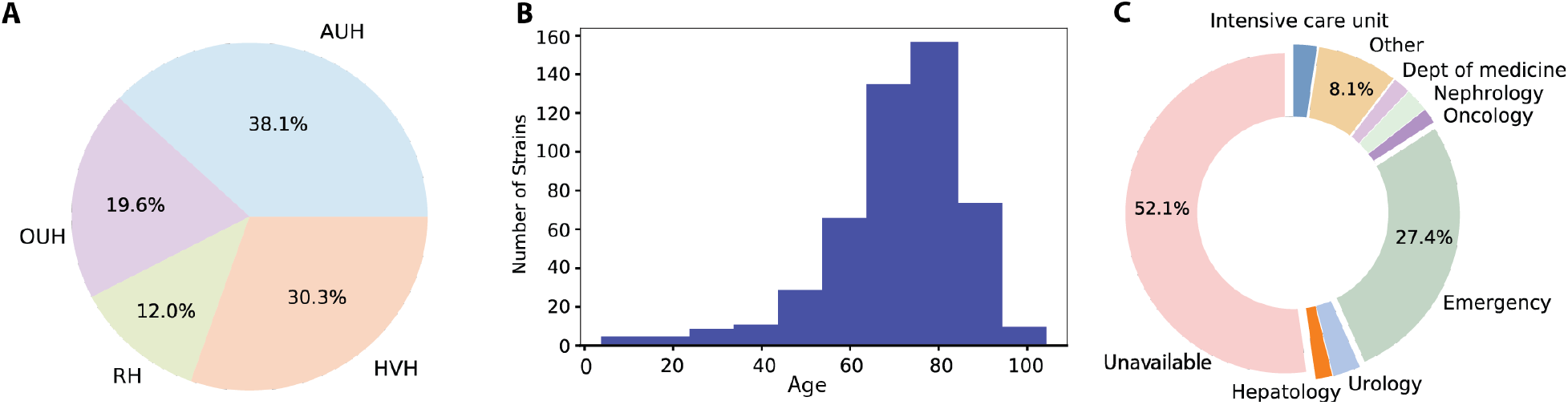
Metadata of 419 Klebsiella pneumoniae (KP) strains collected in Denmark. A) KP strains were collected from four Danish hospitals. OUH = Odense University Hospital, AUH = Aarhus University Hospital, HVH = Hvidovre Hospital, RH = Rigshospitalet. 2) Age distribution of patients where the KP strains were isolated from. 3) Hospital departments where strains were isolated.

To contextualize these strains in relation to KP strains isolated around the world, we also downloaded a set of 2,124 KP genome sequences from PATRIC that were also isolated from humans with accompanying AMR profiles (see methods). These strains were collected from 13 countries in North America, South America, Europe, Asia and Oceania, with 76% of the strains isolated from the United States, 9.7% from the United Kingdom, 4.6% from Australia, 4.5% from Thailand and 3% from Nepal (Fig. S1). This set of diverse strains enables the comparison in characteristics and antibiotic resistant properties between Danish isolates collected in this study and strains from other geographical areas.

### General characteristics of genome sequences

Genome sequences were generated to enable a comprehensive evaluation of the Danish clinical KP strains. The collected isolates were sequenced using Illumina HiSeq2000 with 150bp pair-end reads, assembled with Unicycler v1.2 and annotated using Prokka v1.13. The N50 scores of these 491 genome sequences range between 100K to 743K, with an average of 296.7K. The number of coding genes in each genome sequence ranges between 4,387 and 5,491, which is a range observed for other KP strains. We built a phylogenetic tree of the 491 Danish strains using Phylophlan and found that strains from the same hospitals did not cluster together, suggesting that this set of strains does not contain isolates that were transmitted within the hospital.

The genome sequences were further characterized using Kleborate (Lam, Wick, et al. 2018; Lam, Wyres, et al. 2018; Wyres et al. 2016; Wick et al. 2018), a software developed to screen KP genome sequences for their species, sequence types (STs) and other properties using known knowledge bases. Kleborate analysis suggests that while all strains belong to the KP species complex that is composed of seven species, not all of them belong to the KP species. Out of all 491 strains, only 412 are KP strains, while the remaining 16.1% of the strains belong to *K. quasipneumoniae subsp similipneumoniae, K. quasipneumoniae subsp quasipneumoniae and K. variicola subsp variicola*. The majority of strains (96.8%) in PATRIC dataset belong to *K. pneumoniae* (Fig. 2A). Kleborate also reported that these Danish strains have very diverse sequencing types - 247 sequence types exist across Danish strains with 10.2% strains belonging to ST391, 5.9% strains belonging to ST37, 2.9% belonging to ST20 and 2.4% belonging to ST14, respectively (Fig. 2A). In comparison, PATRIC strains are more concentrated in specific STs - around 25% PATRIC strains belong to ST258 and 21.5% belong to ST307 (Fig. 2C). Both STs are well studied drug resistant groups and causes of hospital-associated infections, and they are both underrepresented in the Danish strains - only two strains were predicted to belong to ST307, and no strains belonging to ST258 are present. Since more than 76% of the PATRIC strains were isolated from the United States, our observation is consistent with previous studies that suggest ST258 likely emerged and remains prevalent in the United States (Wyres and Holt 2016).

**Figure 2:**
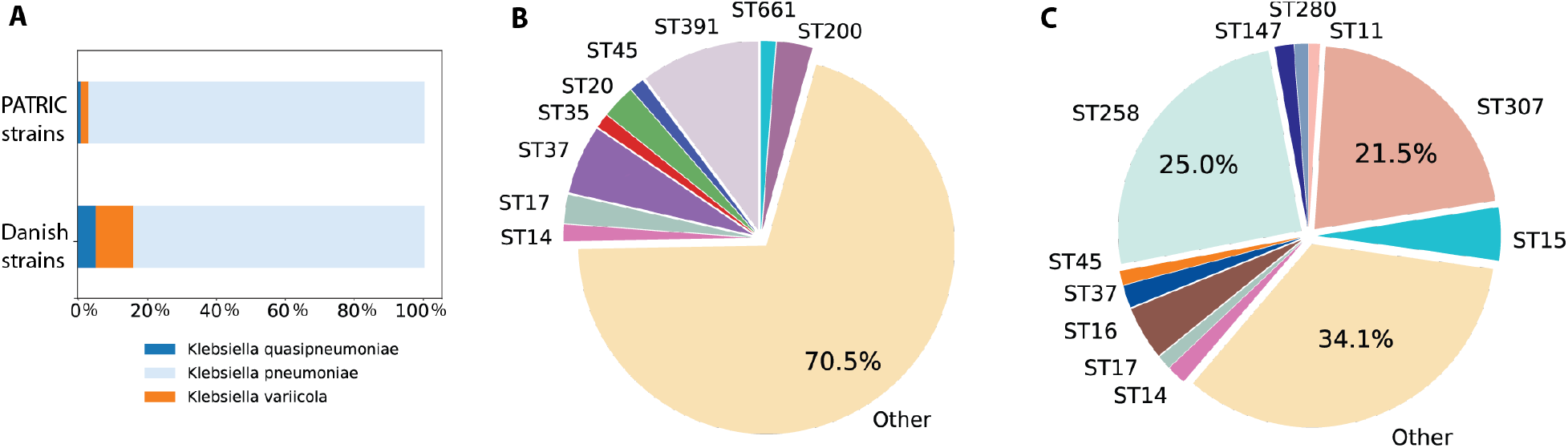
Species and sequencing types. A) Species of strains in Danish and PATRIC datasets B) Sequencing type distribution of Danish strains. C) Sequencing type distribution of PATRIC strains.

We also performed an analysis to identify plasmids in these isolates as plasmids are heavily involved in transfer of virulence factors and AMR genes across strains and species. The genome sequences of 491 Danish strains were mapped to the PlasmidFinder database (Carattoli et al. 2014) using BLAST, and plasmids with more than 80% sequence similarity and greater than 80% alignment length were considered present. Out of all strains, 362 strains have at least one plasmid identified (Fig. S2) and a total of 51 unique plasmids were identified. While most plasmids were only present in a few strains, plasmid FIB(Kpn3) was identified in 303 strains (61.7%). It encodes genes that confer resistance to toxic compounds, metals and antimicrobials, as well as an iron(III) uptake system (García-Fernández et al. 2012). Additionally, the following six plasmids seem to be present in the same subset of more than 100 strains: IncFII(K), IncFII(S), IncFII(Yp), IncFII(p14), IncFII(pRSB107), IncFII_1_pKP91 that includes various plasmid virulence involving fimbriae, capsular, aerobactin, genes conferring resistance to metals and multiple antibiotics. (Villa et al. 2010; Carattoli et al. 2014). The prevalence of plasmids does not seem to be correlated to which hospital the strain was isolated from (Fig. S2), suggesting the potential transmission of these plasmids across Denmark, rather than just certain hospitals or regions.

### Genomic content varies significantly across strains

A pangenome of KP was constructed to evaluate shared and unique gene content across the strains. The pan-genome is defined as all genes present in the strains of a species, which is composed of core genome (genes shared by all strains), accessory genome (genes present in some strains) and unique genome (genes only present in one strain). We first constructed a pan-genome for the Danish strains alone using the 491 annotated genomes by grouping similar genes (sequence similarity > 80%) into the same gene cluster using CD-HIT. There are in total 27,412 gene clusters, of which 2,560 are core gene clusters shared by all 491 strains, and 8,293 unique gene clusters that are only present in one strain (Fig. 3A). We then incorporated the PATRIC strains to build a much larger and comprehensive pan-genome. The pan-genome constructed suggest that the KP strains have significant variations in genomic content across strains. The addition of 2,124 PATRIC strains added another 14,816 gene clusters and shrank the core genome shared by all 2,615 strains to 1,559 genes (Fig. 3A). The pan-genome plot suggests that KP strains likely have an open pangenome, that is, each additional genome will add new gene content to the pan-genome (Fig. 3B), which is consistent with previous literature findings (Holt et al. 2015).

**Figure 3.**
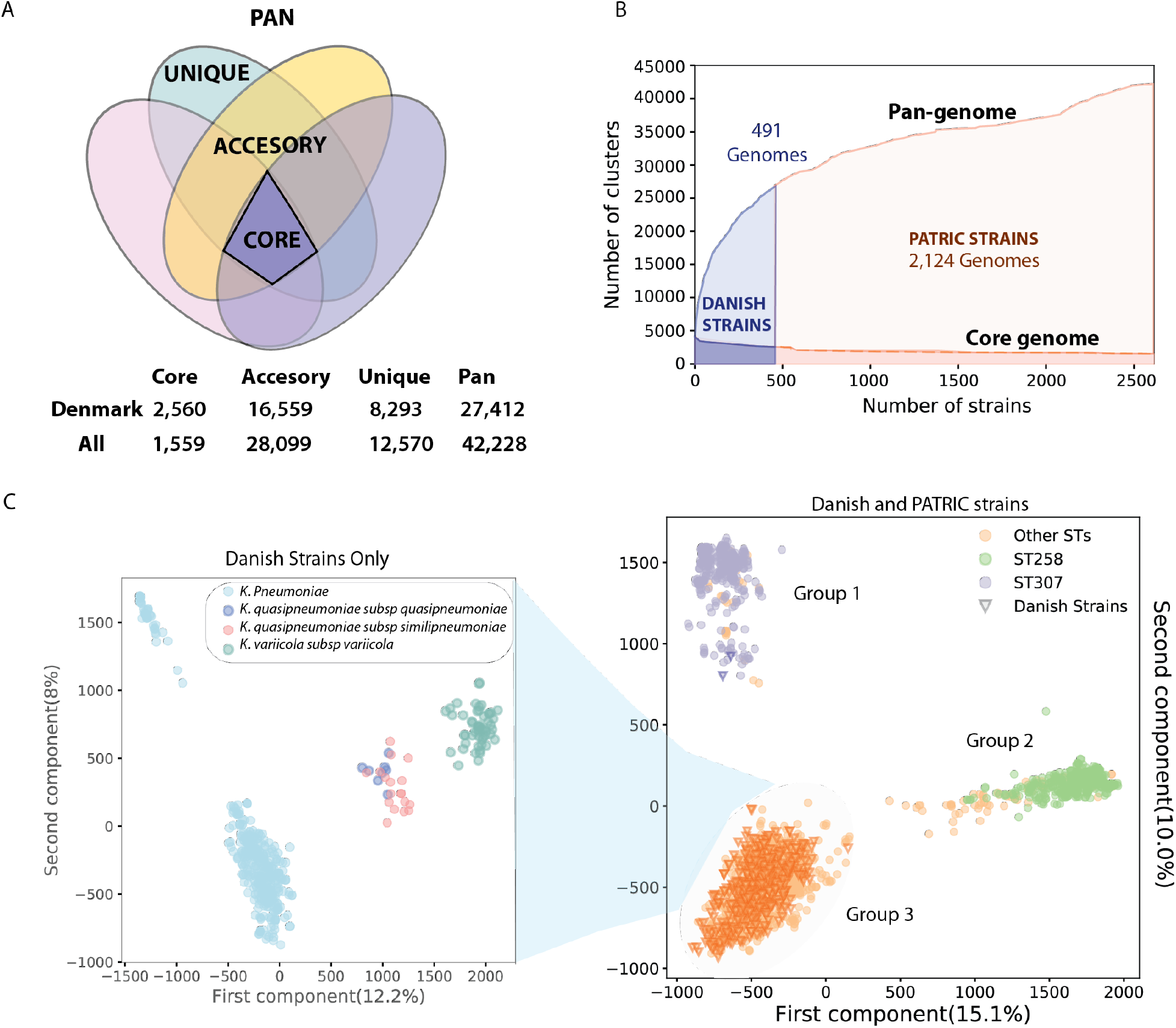
Pan-genome analysis of Danish and PATRIC strains. A) Number of core, accessory and unique genes. B) Changes in pan-genome and core genome for each additional strain, broken down into Danish and PATRIC strains. C) Plot of first two principal components in PCA analysis of pan-genome. Plot on the left is focused on Danish strains only, while plot on the right includes strains in both datasets.

### Separation by sequencing types is related to AMR genes

The principle component analysis (PCA) on the pan-genome suggests subgroups exist within this set of 2,615 strains due to differences in genomic content. The first and second principle components explained 15.1% and 10% of the variance in the pan-genome, respectively. Visualization of the first two principal components suggest three potential subgroups within the entire set of strains (Fig. 3C right). Interestingly, the three groups are separated based on sequence types, rather than species, as group 1 is represented by ST307 and similar STs, group 2 is represented by almost all ST258 strains and few other similar STs, suggesting that the genomic differences across STs are even bigger than that across different species. The majority of Danish strains fell in group 3, due to the lack of ST307 and ST258 and other similar STs. Interestingly, the set of Australian strains all fell within group 3 as well, while other countries had strains falling into all 3 groups. Specifically, the majority of the strains from ST258 and ST307 came from the United States. This result suggests potential differences in the gene portfolios of KP strains across different countries, which could be a result of different antimicrobial usage guidelines in different countries and import and spread of KP AMR clones (Klein et al. 2018).

Many gene clusters that separated ST307, ST258 and similar STs from the other strains are involved in AMR. We identified the top 100 gene clusters in PC1 and PC2 respectively (Table S1) and looked into their distribution in all strains. The result suggested that many differentiating gene clusters were only uniquely present in the ST307 group or ST258 group. More detailed analysis revealed that while most of the gene clusters are hypothetical proteins with no available annotations, some known genes are involved in antibiotic resistance. In PC1 that separates ST258 group and the other strains, the top loading genes include Tyrosine recombinase *XerC* (Cameranesi et al. 2018), Prophage integrase IntA and antirestriction protein *KlcA* (Liang et al. 2017) that are known to disseminate AMR genes, efflux pump membrane transporter *BepE* and Toluene efflux pump periplasmic linker protein *TtgG* that can export drugs, and toxic protein *SymE*, a DNA damage-inducible protein involved in repairing DNA and RNA damage. These genes are logical because ST258 is one of the most studied drug resistant STs. While the rest of the gene clusters may not have functional annotation or direct evidence related to AMR, they can still potentially contribute to acquisition of AMR (via increased fitness or other mechanisms) and are worth pursuing in future experiments for verification (Table S1). In PC2 that separated ST307 and other strains, some top loading genes clusters also contribute to AMR, such as multiple restriction enzymes, Tellurite resistance protein *TehA*, DNA replication and repair protein *RecF*, but most other annotated gene clusters are involved in metabolic functions, such as metabolism of UDP-glucose, fumarate, L(+)-tartrate and Alpha-maltose-1-phosphate. A previous study focused on the strain-specific models of KP also revealed the potential linkage between AMR and metabolic functions (Norsigian et al. 2019).

PCA analysis focused on Danish strains only revealed subgroups due to gene content in different species. We identified four potential subgroups within the Danish strains (Fig. 3C left) that are separated based on their species, - *K. quasipneumoniae and K. variicola* each form a group, but two groups exist in KP species. Further analysis suggests that the separation within the KP groups is due to the differences across sequence types, as ST391 which formed the upper left cluster was shown to have large variation in metabolic genes compared to other STs in this dataset, and therefore formed a separate group in the PCA analysis.

### Danish strains contain fewer virulence factors/genes than strains from other countries

To perform a comprehensive evaluation of the virulence factors found across the KP isolates, we identified virulence factors in Danish strains using the curated VF database VFDB. We mapped the genome sequences of the KP strains to the nucleotide sequences of curated VFs in VFDB using BLAST, and VFs with more than 80% sequence similarity and >80% alignment lengths are considered present in the strain. One hundred and sixty eight VFs were found present in at least one strain, including VFs involving secretion systems, capsule/ lipopolysaccharide biosynthesis, iron sequestration, adhesion, and more (see Table S2 for details). For PATRIC strains, 242 VFs were found to be present in at least one strain.

Specifically, we focused on some of the well-studied virulence loci in KP - the four key acquired virulence factors that contributes to hypervirulence, including siderophore yersiniabactin (*ybt*), aerobactin (iuc), salmochelin (*iro*), and the genotoxin colibactin (*clb*). In particular, a set of 20 VFs involved in biosynthesis and regulation of yersiniabactin are only present in around 32.6% strains that came from different hospitals. Yersinibactin is an iron uptake system widely spread in pathogenic bacteria, enabling these bacteria to acquire iron for essential cellular processes. Salmochelin and aerobactin were originally found in *Salmonella* and *Escherichia coli* and have similar functions in sequestering iron. The gene clusters responsible for synthesis and functions of salmochelin (*iroBCDEN*) and aerobactin (*iucABCD*) were found in only 16 and 17 strains that largely overlap with each other. Additionally, 18 VFs involved in colibactin biosynthesis which can cause DNA double strand breaks in human cells are only present in a subset of 9 strains. These strains with additional colibactin gene clusters would likely pose higher risk for their host, as colibactin is known to promote colorectal tumorigenesis (Faïs et al. 2018). We also examined two hypermucoidy genes *rmpA* and *rmpA2*, which are known as the “regulators of mucoid phenotype” and can up-regulate capsule production (Holt et al. 2015). We identified 15 strains with *rmpA* and 8 strains with *rmpA2*. Kleborate was also used to calculate the virulence score using the above mentioned four key VFs - the Kleborate reports that the average virulence score is 0.15 (on a scale of 0-5) with a standard deviation of 0.85 for Danish strains. For the PATRIC strains however, the average reported virulence score is around 1.17 with a standard deviation of 0.76, as they have more acquired virulence factors compared to the Danish isolates. Among them, strains from the United states have an even higher virulence score of 1.26.

### Danish strains appear to be less antibiotic resistant in experiments

In total, twenty antibiotics and antibiotics combinations were tested for the Danish strains using the disk diffusion method and AMR phenotype was determined according to the EUCAST guidelines. Because the AMR profiles were generated separately by individual hospitals where the strain was collected, not all strains were tested for the same antibiotics. As shown in the figure, only 6 out of 20 antibiotics (Cefuroxime, Ciprofloxacin, Gentamicin, Meropenem, Pivmecilinam and Piperacillin/Tazobactam), were tested for all strains, while the rest of the antibiotics were tested only on a subset of the strains. For all 20 antibiotics tested, 15 antibiotics had resistance rates lower than 20%. All strains are shown to be resistant to Ampicillin as expected due to the presence of the SHV beta-lactamase in the core genome (Wyres and Holt 2016). In addition, a subset of Danish strains were resistant to Ceftolozane/Tazobactam (31.6%), Trimethoprim (31.6%), Pivmecillinam (23.2%) and Trimethoprim/Sulfamethoxazole (20.7%).

The PATRIC strains seem to be more resistant to antibiotics. In total, AMR profiles were generated for 57 antibiotics due to the large number of strains involved, of which 16 antibiotics were also tested in the Danish strains. The AMR profiles of PATRIC strains were generated using various methods and standards (CLSI and EUCAST). For comparison, we calculated the percentage of strains that are resistant to each antibiotic for these 16 antibiotics and compared them across the two datasets. Besides Ampicillin, the resistance rate in PATRIC dataset is higher than that of Danish strains for all antibiotics tested, and for half of the antibiotics, the resistance rate of PATRIC strains are more than 5 times that of Danish strains (Fig. S3).

### Less known AMR determinants identified in Danish strains

We screened AMR genes and allelic variants against the ARG-Annot database using Kleborate and the reported AMR determinants are broken down into 19 drug classes. For Danish strains, the results suggest that at least one strain is potentially resistant to 15 out of 19 classes of drugs, with prevalent resistance predicted in β-lactam antibiotics, resistance to aminoglycosides, sulfonamides and trimethoprim in a subset of strains, and limited resistance to tetracyclines, fluoroquinolones, and phenicols in few strains. The presence of AMR determinants does not seem to correlate with hospitals, and comparison with PATRIC strains suggest that the prevalence of AMR determinants in Danish strains is significantly lower (Fig. S4). Kleborate predicts that on average Danish strains are resistant to 1.56 classes of drugs, with 159 strains not resistant to any antibiotics and 203 strains resistant to 1 class of drugs. PATRIC strains were predicted to be in general much more drug resistant, as they were predicted to be resistant to 5.7 classes of drugs on average. This is consistent with the phenotypic results discussed in the last section.

We also examined extended-spectrum beta lactamases (ESBL) and carbapenem resistance genes as they are both of particular clinical concerns (Wyres and Holt 2016). Consistent with experimental results on carbapenem (meropenem and imipenem) (Fig. 4A), very few carbapenemse-producing strains were detected. We only found three strains that possess the carbapenem resistance genes blaOXA-48 and one other strain containing blaOXA-232. ESBL genes are slightly more prevalent in these strains, 14.1% (Fig. 4B), therefore resulting in higher resistance in drugs belonging to cephalosporins including ceftazidime, cefuroxime, cefpodoxime and ceftriaxone (Fig. 4A). A total of nine variants of two ESBL genes *blaSHV* (SHV-13,SHV-12,SHV-101,SHV-27,SHV-41,SHV-42) and *blaCTX-M* (CTX-M-15, CTX-M-14, CTX-M-3) were identified in 69 strains, and each strain only possessed one ESBL gene (Table S3).

**Figure 4.**
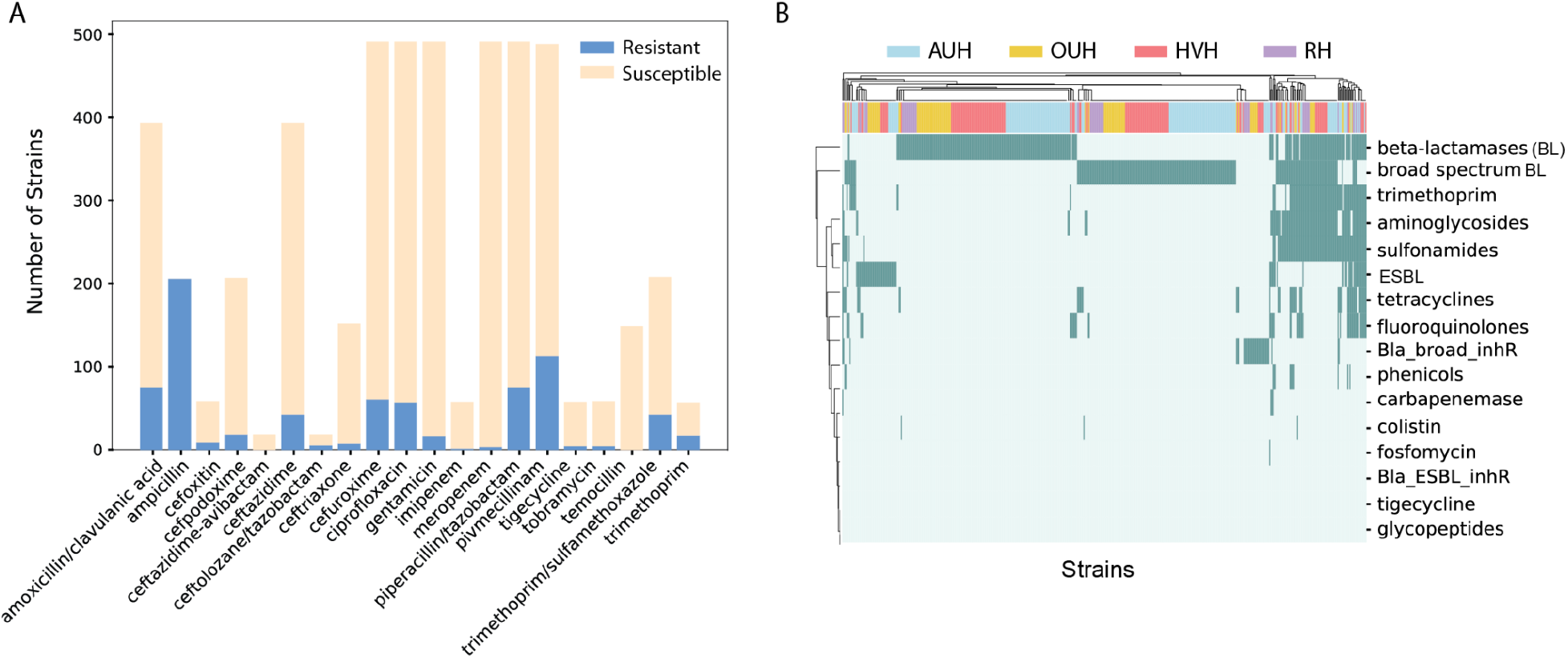
Antibiotic resistance of Danish strains. A) AMR profiles for 20 drugs were experimentally tested. B) Detection of antibiotic determinants in genomes sequences, labeled by hospitals where the strains were isolated.

### Discovery of potential AMR determinants

Comparison between experimental data and predicted resistance based on known AMR determinants suggests that for most antibiotics, presence of AMR determinants can be used to predict AMR resistance accurately, but there still exist some inconsistencies that are potentially knowledge gaps. For example, for aminoglycosides, around 18.7% of the strains were predicted to have AMR determinants including *StrA, StrB, Aac, AadA1-pm*, but phenotypic data for both antibiotics belonging to aminoglycosides showed comparatively lower resistance: Gentamicin (3.5%) and tobramycin (8.5%). Further analysis of aminoglycosides determinants suggest that while *StrA* and *StrB* encodes aminoglycoside-3”-phosphotransferase(APH(3”)-Ib) and aminoglycoside-6-phosphotransferase (APH(6)-Id) (Chiou and Jones 1995) that confer resistance to streptomycin, an antibiotic that also belongs to aminoglycoside (Vakulenko and Mobashery 2003), it does not necessarily confer resistance to gentamicin and tobramycin, as enzymes encoded by *StrA* and *StrB* are likely specific to streptomycin.

There were also cases where we observed higher resistance phenotypically than that predicted based on presence of known AMR genes, which could be an opportunity for potential discovery of new AMR determinants and drug targets. One example is tigecycline, an antibiotic derived from tetracycline designed to overcome mechanisms of tetracycline resistance (Pournaras et al. 2016). No tigecycline-related determinants were reported in any strains in the genomic analysis, yet 8% percent of the tested Danish strains were shown to confer resistance (Fig. 4), suggesting there may be unknown tigecycline-related determinants that were not reported. We also observed similar results in PATRIC strains, while no AMR determinants were reported, around 37% of the tested strains conferred resistance phenotypically.

To uncover potential unknown genes conferring resistance to tigecycline, we performed genome-wide association study (GWAS) using the previously constructed pan-genome. Because tigecycline was only tested in a sub-set in one of the four hospitals (RH), data was only available for 58 Danish strains. To enhance the power of our analysis, we included another 131 PATRIC strains with tigecycline resistance data. For all 189 strains, we performed GWAS analysis using both analysis of variance (ANOVA) and chi-squared with the pan-genome matrix and tigecycline resistance profile determined experimentally. We found the top seven genes identified by chi-squared can best differentiate the resistance and susceptible strains (Table 1, Fig. S5) and these genes were also among the top genes identified by ANOVA.

**Table 1:** The top seven genes identified that best differentiate the strains that are resistant and susceptible to tigecycline.

| Rank | Gene Name | Function | % resistant if present | % resistant if absent |
| --- | --- | --- | --- | --- |
| 1 | Unknown | IS1380-like element ISEc9 family transposase | 72.8% | 15.2% |
| 2 | *tetR* | Tetracycline repressor protein class A from transposon 1721 | 77.1% | 17.5% |
| 3 | *tetA* | Tetracycline resistance protein, class C | 76.5% | 18.1% |
| 4 | *yedA* | Predicted inner-membrane transporter | 76.5% | 18.1% |
| 5 | Unknown | Unknown | 67.4% | 16.1% |
| 6 | *bla* | Beta-lactamase CTX-M-1 | 66.7% | 16.7% |
| 7 | *aer* | Aerotaxis receptor | 69.7% | 19.9% |

Interestingly, genes involved in tetracycline resistance were among the top genes that potentially contribute tigecycline resistance. *tetA* encodes an active tetracycline efflux and tetR is a repressor of tetA. Binding of the tetracycline to tetR will reduce the affinity of tetR to tetA promoter sites to enable resistance to tetracycline. Out of all 189 strains tested for tigecycline, all strains have tetA (class B) but only 34 strains have tetA (class C) that share 45% sequence identity with tetA(class B) (Yin et al. 2000). tetR is also present in the same 34 strains plus one other strain. Although tigecycline is to be expected to overcome resistance mechanisms of tetracycline, previous studies have shown that the presence of tetA or mutations in tetA is associated with tigecycline resistance (Ahn et al. 2016; Du et al. 2018). The top determinant is similar to a ISEc9 family transposase family transposase, an insertion sequence known to be associated with insertion of ESBL genes into KP (de Man et al. 2018). We also identified a predicted transporter encoded by yedA, which was also identified as a candidate resistance gene for doxycycline (a tetracycline antibiotic) in a previous study (Mahfouz et al. 2018). It is interesting that bla showed up in the selected genes as tigecycline does not belong to betalactam antibiotics, but previous studies showed that Enterobacteriaceae carrying bla_CTX-M_ confer moderate resistance against tigecycline (Zeynudin et al. 2018; Ahn et al. 2016). Little information can be found on the hypothetical protein, aerotaxis receptor and their relationship with AMR, but may be potential opportunities for future exploration.

In addition to gene-level analysis, we have also performed GWAS on the variants of all gene clusters and identified two variants of interest. We first identified a truncated variant (with only the last 45 amino acids in the protein) of elongation factor Tu encoded by gene tufA that is associated with tigecycline resistance. Out of 27 strains of this tufA variant, 22 confer resistance to tigecycline. It is interesting as tufA promotes the GTP-dependent binding of aminoacyl-tRNA to the A-site of ribosomes during protein biosynthesis (Caldas, El Yaagoubi, and Richarme 1998), while tigecycline blocks the interaction of aminoacyl-tRNA with the A site of the ribosome to inhibit protein synthesis (Slover, Rodvold, and Danziger 2007). However, from literature search it is unclear if this variant of tufA can bypass the tigecycline inhibition, or interfere with tigecycline mechanism in any way, which may be better explained by future experiments. One other variant of gene tnpR was also identified that encodes Transposon Tn3 resolvase, which has a point mutation in the third amino acid (Isoleucine to Leucine). Twenty-eight out of 38 strains with this variant presented tigecycline resistance. Tn3 often carries beta-lactamase but the role of this mutation of Tn3 resolvase is yet to be further explored.

Due to the limited sample size, attempts to select novel determinants of tigecycline using machine learning methods were unsuccessful. However, such efforts have proven to be productive when the datasets are large enough, as it is able to incorporate interactions between genes compared to univariate tests as we did in the GWAS analysis.

## DISCUSSION

Through collection and analysis of 491 Danish KP strains, we were able to obtain a deeper understanding of KP strains in Denmark that may contribute to development of treatment: 1) Great genomic diversity exists in KP strains, with a total of 27,412 gene clusters identified in 491 Danish strains. 2) Danish strains differ from PATRIC strains collected from other countries, likely due to the difference in the lineage. 3) Danish strains appear to be less antibiotic resistant and contain fewer virulence genes. 4) Potential AMR determinants were identified for tigecycline through GWAS analysis.

Some observations from our study are consistent with the previous understanding of KP. The majority of KP strains belonged to the species *K. pneumoniae*, as *K. varricola* and *K. quasipneumoniae* were known to be less virulent (Long, Linson, et al. 2017). As shown by our and previous studies (Holt et al. 2015), KP has an open pan-genome, with many AMR and virulent genes mobilized by a variety of plasmids and other conjugative elements (Lam, Wick, et al. 2018; Ramirez et al. 2014). It was also not surprising to see the majority of the KP strains isolated from older patients, as these patients tend to have weaker immune systems. However, while many previous studies suggest KP can cause within-hospital transmission (Gu et al. 2018; Snitkin et al. 2012), we did not see such signs in our analysis.

Danish strains differ from strains from other countries in several aspects. First, while Danish strains have a range of STs, two most studied AMR STs - ST258 and ST307 were not present in this collection of strains, which results in a genomic content difference between Danish strains and PATRIC strains from other countries as shown in Fig 3C. Our analysis suggested that many genes that differentiate these two STs from Danish strains are involved in AMR. While both ST258 and ST307 were reported to have spread globally (Villa et al. 2017; Bowers et al. 2015), they may not yet have become endemic in Denmark, which could explain the relatively low AMR and virulence scores reported in the Danish strains. Nonetheless, high rates of recombination within KP and the fluid dynamics of gene flow within KP populations (Comandatore et al. 2019) create risk that these lineages (or the genetic factors underlying them) could become prevalent in Denmark in the future.

Although the AMR experimental results mostly matched the genomic prediction based on AMR gene determinants, we were able to propose potential AMR determinants for tigecycline where predicted AMR is lower than experimental measurements. Through GWAS analysis, we proposed 7 genes and 2 variants that could potentially contribute to tigecycline resistance, of which some results also overlapped with findings in previous studies.

It is important to note certain limitations of our study. First, we have used PATRIC strains with AMR profiles as a comparison to our dataset, but the PATRIC database could be biased towards well-studied AMR sequencing types, which can potentially contribute to the relatively high AMR and virulent scores of PATRIC strains. Second, only a very limited amount of metadata was collected from patients, which undermines our ability to perform more in-depth analysis. Lastly, different antibiotics were tested in each hospital, which reduced the sample size for certain antibiotics and limited the statistical power of the analysis. Despite the limitations, our study still provided valuable information on the population structure, virulence level and antibiotic resistance of KP strains in Denmark. This study motivates similar efforts to collect clinical isolates from a larger population, which will likely provide more information to understand and combat ARM in KP.

## METHODS

### Strain collection

The 491 *Klebsiella Pneumoniae* (KP) strains were collected from positive blood culture isolates from clinical microbiology departments of four Danish hospitals (Aarhus University Hospital, Hvidovre Hospital, Odense University Hospital, and Rigshospitalet) between January 2018 and December 2018. These hospitals have an uptake area covering 51% of the Danish population.

### Ethical Consideration

The local ethics committee at the Capital Region of Denmark (Region Hovedstaden) approved the use of the clinical Klebsiella spp. isolates: registration number H-19029688

### Antibiotic resistance measurement

Antibiotic susceptibility testing was done on Müller-Hinton agar plates (State Serum Institute, Diagnostica A/S, Hillerød, Denmark) using 0.5 McFarlan according to the European Committee on Antimicrobial Susceptibility Testing (EUCAST) guidelines (ESCMID-European Society of Clinical Microbiology and Infectious Diseases n.d.)

### Genome sequence generation

Genomic DNA was prepared from pure culture of Klebsiella isolates using DNeasy Blood and Tissue kit (Qiagen) and whole genomes were sequenced on an Illumina HiSeq 2000 platform. All isolates were sequenced using 2×150-bp paired-end sequencing to obtain an average of 80X genomic coverage.

The paired end reads are filtered and assembled with Unicycler v1.2. We filtered out genome assemblies with N50 scores less than 100,000. The N50 scores of the remaining 491 genome sequences range between 100,000 to 743,000, with an average of 296,700. The assemblies are then annotated using Prokka v1.13. The number of coding genes in each genome sequence ranges between 4,387 and 5,49.

PATRIC strains were downloaded from the public database PATRIC. We downloaded all genome sequences of *Klebsiella Pneumoniae* strains with accompanying AMR data as of September 2019. We applied the same filtering criteria to these strains and filtered out strains with N50 scores less than 100,000 and ended up with 2,124 strains. We also annotated these strains Prokka v1.13 to ensure consistency with Danish strains.

### Genome sequence characterization

Species and serotypes were characterized using the software tool Kleborate following the instructions (https://github.com/katholt/Kleborate). Species identification is accomplished through comparison with known *Klebsiella* genomes, and serotype is determined through multilocus sequence typing using the 7-locus scheme.

The plasmids were identified through mapping the assembled genomes to the PlasmidFinder database using BLAST. Plasmids with more than 80% similarity and more than 80% alignment length were considered to be present in the genome.

### Pan-genome analysis

The annotated genome sequences were aligned against each other using CD-HIT, with the cutoff for “align average” set to 80% so that genes with more than 80% sequence similarity were grouped together as one gene cluster. A binary pan-genome matrix is then generated based on the presence and absence of gene clusters, and core, accessory and unique gene clusters were identified. PCA was performed on the pan-genome matrix using the PCA function in Sklearn. Data points were plotted for the first two principal components, and metadata (including country, serotype, species and hospitals) was used to label the data points to further understand the PCA plot. We also investigated the important genes in the first two principal components by examining the gene clusters with the highest absolute values of their coefficients in PC1 and PC2.

### Virulence factors characterization

Virulence factors were identified by mapping the genome sequences to the curated database VFDB using BLAST. VF with more than 80% sequence similarity and alignment length are considered present in the genome sequence. We specifically looked into the key acquired VFs in KP, including *ybt, iuc, iro* and *clb*, as well as the two genes *rmpA* and *rmpA2* responsible for hypermucoidy. Kleborate also reported results on the above key VFs and assigned VF scores to the strains based on the presence/absence of the VFs.

### AMR determinants identification

Kleborate was used to identify AMR determinants in the genome sequences using the -- resistance option. Kleborate identified known AMR determinants by genome sequences to the ARG-Annot database of the acquired resistance genes, and reported the best matching variants. The AMR determinants were also grouped by the drug class they confer resistance to according to the ARG-Annot, with beta-lactamases broken down into Lahey classes. Kleborates also reports the predicted number of drug classes each strains is resistant to based on the presence of AMR determinants.

